# The discovery of a recombinant SARS2-like CoV strain provides insights into SARS and COVID-19 pandemics

**DOI:** 10.1101/2020.07.22.213926

**Authors:** Xin Li, Xiufeng Jin, Shunmei Chen, Liangge Wang, Tung On Yau, Jianyi Yang, Zhangyong Hong, Jishou Ruan, Guangyou Duan, Shan Gao

## Abstract

In December 2019, the world awoke to a new zoonotic strain of coronavirus named severe acute respiratory syndrome coronavirus-2 (SARS-CoV-2). In the present study, we identified key recombination regions and mutation sites cross the SARS-CoV-2, SARS-CoV and SARS-like CoV clusters of betacoronavirus subgroup B. Based on the analysis of these recombination events, we proposed that the Spike protein of SARS-CoV-2 may have more than one specific receptor for its function. In addition, we reported—for the first time—a recombination event of *ORF8* at the whole-gene level in a bat and ultimately determined that ORF8 enhances the viral replication. In conjunction with our previous discoveries, we found that receptor binding abilities, junction furin cleavage sites (FCSs), strong first ribosome binding sites (RBSs) and enhanced *ORF8*s are main factors contributing to transmission, virulence and host adaptability of CoVs. Junction FCSs and enhanced *ORF8*s increase the efficiencies in viral entry into cells and replication, respectively while strong first RBSs enhance the translational initiation. The strong recombination ability of CoVs integrated these factors to generate multiple recombinant strains, two of which evolved into SARS-CoV and SARS-CoV-2 by nature selection, resulting in the SARS and COVID-19 pandemics.

## Introduction

A new zoonotic strain of coronavirus named severe acute respiratory syndrome coronavirus-2 (SARS-CoV-2) emerged in December 2019. Since SARS-CoV-2 is high similar to SARS-CoV, many studies focused on the investigate of the receptor binding domain (RBD) of the Spike protein and its receptor ACE2 using the same strategies and methods as in SARS-CoV [1]. Different from these studies, we previously reported several other findings on SARS-CoV-2 for the first time, including the following in particular: (1) the alternative translation of Nankai coding sequence (CDS) that characterize the rapid mutation rate of betacoronavirus at the nucleotide level [2]; (2) a furin cleavage site (FCS) “RRA**<u>R</u>**” in the junction region between S1 and S2 subunits (junction FCS) of SARS-CoV-2 that may increase the efficiency of viral entry into cells [3]; and (3) the use of 5’ untranslated-region (UTR) barcoding for the detection, identification, classification and phylogenetic analysis of—though not limited to—CoVs [4]. By data mining betacoronaviruses from public databases, we found that more than 50 nucleotides (nts) at the 3’ ends of the 5’ UTRs in betacoronavirus genomes are highly conserved with very few single nucleotide polymorphisms (SNPs) within each subgroup of betacoronaviruses. We defined 13~15-bp sequences of 5’ UTRs including the start codons (ATGs) of the first open reading frames (ORFs) as barcodes to represent betacoronaviruses. Using 5’ UTR barcodes, 1265 betacoronaviruses were clustered into four classes, matching the C, B, A and D subgroups of betacoronavirus [4], respectively. The 3’ end of each 5’ UTR includes the first ribosome binding site (RBS) of betacoronavirus, which regulates the translational initiation of downstream genes ORF1a and 1b [5]. In particular, SARS-CoV-2 and SARS-CoV have the strong first RBSs enhancing the translational initiation [4]. These previous studies indicated that receptor binding abilities, junction FCSs and strong first RBSs are main factors contributing to transmission, virulence and host adaptability of CoV.

The present study started with the identification of key recombination regions and mutation sites cross three clusters of betacoronavirus subgroup B. Using the insertions and deletions (InDels) at six sites, we identified two recently detected betacoronavirus strains RmYN01 and RmYN02 from a bat [6] and discovered that RmYN02 was a recombinant SARS2-like CoV strain. This led us to report—for the first time—a recombination event in open reading frame 8 (ORF8) at the whole-gene level in a bat, which had been co-infected by two betacoronavirus strains. *ORF8* (**Figure 2**), existing only in betacoronavirus subgroup B, was considered to have played a significant role in adaptation to human hosts following interspecies transmission [7] via the modification of viral replication [8]. Next, we ultimately determined that ORF8 enhances the viral replication, which is another main factor contributing to transmission, virulence and host adaptability of CoV. In the present study, we analyzed these main factors in the context of betacoronavirus evolution (conjoint analysis of phylogeny and molecular [9]) to explain the SARS and COVID-19 pandemics.

**Figure 1.**
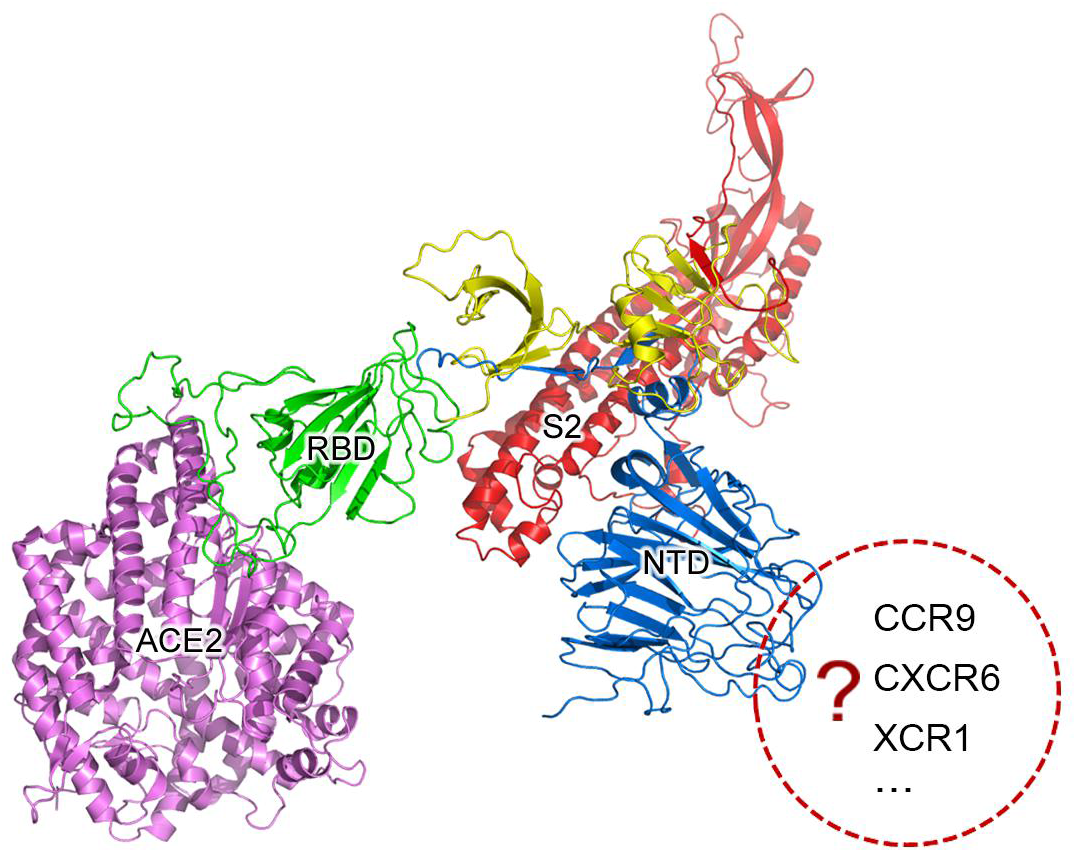
SARS-CoV-2 may have more than one specific receptor. The S protein is cleaved into two subunit S1 and S2 (in red color) for receptor binding and membrane fusion. S1 has two domains, RBD (in green color) and NTD (in blue color). It is well accepted that S1 binds to its specific receptor angiotensin-converting enzyme 2 (ACE2) by the interaction between RBD and ACE2 (in purple color). In the present study, we propose that the Spike protein of SARS-CoV-2 may have more than one specific receptor for its function as gp120 of HIV has CD4 and CCR5. The structure of S was predicted using trRosetta [24].

**Figure 2.**
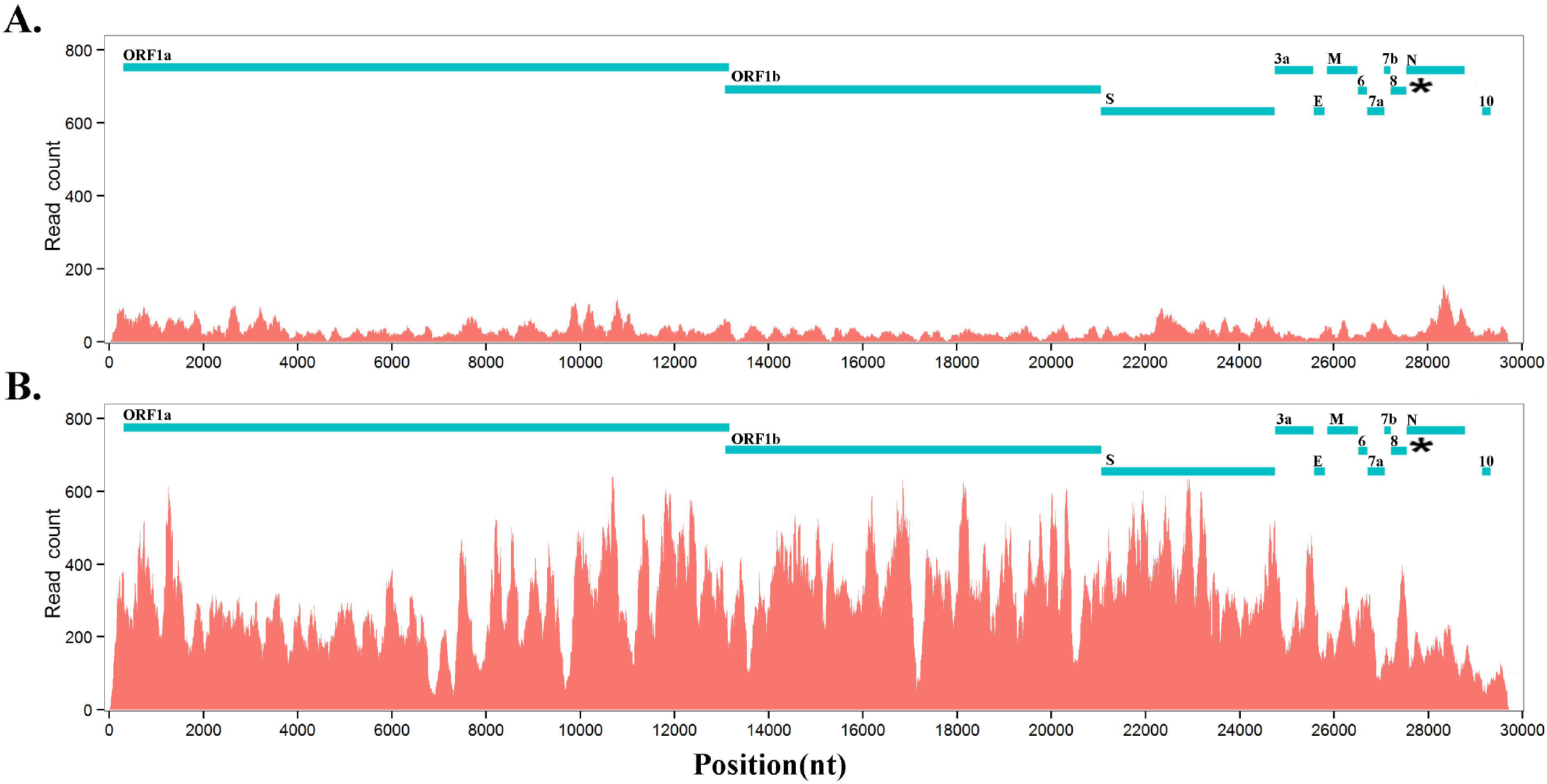
RNA abundances of RmYN01 and RmYN02 in a bat. RNA-seq data from a bat was aligned to two genomes of RmYN01 and RmYN02 (GISAID: EPI_ISL_412976 and EPI_ISL_412977). RNA abundance is represented by read counts (y-axis). The RNA abundance of RmYN02 is 9 times that of RmYN01. **A.** RmYN01 was identified as belonging to the SARS-like CoV cluster and has a type 3 ORF8. **B**. RmYN02 was identified as belonging to the SARS-CoV-2 cluster but has an type 2 ORF8 (enhanced ORF8).

## Results and Discussion

### Identification of key recombination regions and mutation sites

Based on analysis of betacoronavirus subgroup B (**Materials and Methods**), key insertions and deletions (InDels) were identified at six sites (named M1 to M6) in genes *ORF3a*, membrane (*M*), *ORF7a, 7b, 8* and nucleocapsid (*N*), respectively (**Table 1**). Using the InDels at six sites, betacoronavirus subgroup B was classified into two classes: (1) the first class includes SARS-CoV-2 (from humans) and SARS2-like CoV (from animals), and (2) the second class includes SARS-CoV (from humans) and SARS-like CoV (from animals). This classification result is reliable as all recombination and mutations between them are unlikely to undergo reversible changes together. As a mutation site, M1 has a length of 8 nt in the second class and 11 nt in the first class. M2, M3, M4 and M5 in the first class have 3-nt deletions that are complete codons, whereas M6 in the first class has 6-nt deletion that are not complete codons.

**Table 1.**
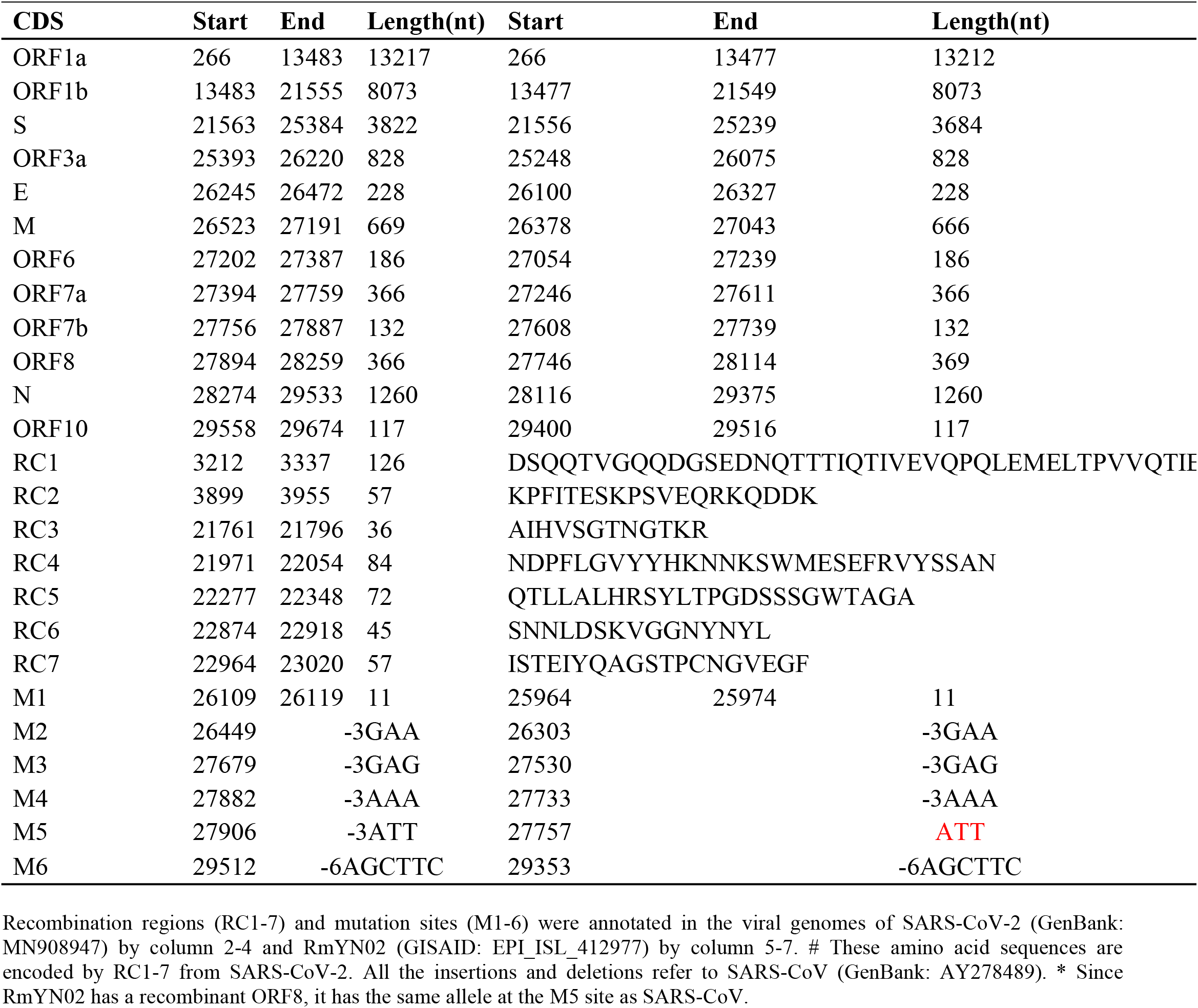
Annotations of recombination regions and mutation sites.

Almost all the identified recombination events occurred in genes *ORF1a, S* and *ORF8* (**Table 1**). The recombination regions RC1 to RC2 and RC3 to RC7 are located in *ORF1a* and the S1 region of the *S* gene, respectively, while the recombination events in *ORF8* are complex (**see below**). To initiate the CoV infection, the S protein encoded by the *S* gene need to be cleaved into the S1 and S2 subunits for receptor binding and membrane fusion. By analysis of all recombination events in all betacoronaviruses, we obtained the following results: (1) there are a few genotypes of each recombination region (RC1 to RC7); (2) RC3 to RC7 have more diversity than RC1 to RC2 in the genotypes; (3) betacoronaviruses within the SARS-CoV-2 and SARS-CoV clusters (**see below**) had the same genotypes of each recombination region; and (4) there were a few non-synonymous substitutions between different sequences of each genotype. These results suggested that recombination, rather than accumulated mutation directly triggered cross-species transmission and outbreaks of SARS-CoV and SARS-CoV-2, while mutation could change potential recombination sites, affecting recombination.

Further analysis showed that two recombination regions (RC6 and RC7) are localized in the receptor binding domain (RBD) of S1 (**Figure 1**), while three other recombination regions (RC3, RC4 and RC5) are localized in the N-terminal domain (NTD) of S1. Almost all the secondary structures of five protein segments encoded by RC3 to RC7 are disordered, which are responsible for protein protein interaction (PPI). This suggested that the recombination of RC3 to RC7 improve the adaptability of betacoronaviruses in new hosts (host range expansion [1]) by enhancing interaction of RBD and NTD with their receptors, exhibiting that the positive selection of S was particularly strong [7]. Since both RBD and NTD had similar recombination events in their PPI regions, we proposed that NTD has a specific receptor as RBD has angiotensin-converting enzyme 2 (ACE2). Thus, the S1 subunit of SARS-CoV-2 may have more than one specific receptor (**Figure 1**) like gp120 of HIV has the cluster of differentiation 4 receptor (CD4) and the CC chemokine receptor 5 (CCR5). Comprehensive analysis and reuse of data from different sources are necessary to determine the other receptor/s of SARS-CoV-2. A previous study identified two genetic susceptibility loci (rs11385942 at locus 3p21.31 and with rs657152 at locus 9q34.2) in COVID-19 patients with respiratory failure using genome-wide association analysis [10]. The locus 3p21.31 was associated to six genes *SLC6A20, LZTFL1, CCR9, FYCO1, CXCR6* and *XCR1*. However, the previous study only focused on the further analysis of the locus 9q34.2 and confirmed a potential involvement of the ABO blood-group system. The researchers did not notice that three chemokine receptors *CCR9*, *CXCR6* and *XCR1* merit further investigation as candidates of SARS-CoV-2 receptors. The analysis of bulk RNA-seq data showed high expression of *CCR9* and *XCR1* in thymus and *CXCR6* in T cells, compared to other tissues and cell types [10]. In particular, the thymic cells were consistently negative for ACE2 and many CoVs can infect thymus [11]. By investigating interaction of three protein segments encoded by RC3 to RC5 in NTD (**Table 1**) with *CCR9, CXCR6* and *XCR1*, we found that *CCR9* is the most possible candidate among three chemokine receptors. However, the final determination of the other receptor/s of SARS-CoV-2 need more calculation and experiments on candidates at the whole-genome level. Our study did not rule out the possibilities of non-receptor proteins binding to NTD.

### Classification of two betacoronavirus strains from a bat

Recently, two betacoronavirus strains RmYN01 and RmYN02 (GISAID: EPI_ISL_412976 and EPI_ISL_412977) were detected from a bat (*Rhinolophus malayanus*) [6]. Since betacoronaviruses from subgroup B share many highly similar regions in their genome sequences, it is very difficult to assemble them correctly using high-throughput sequencing (HTS) data from one sample. Therefore, EPI_ISL_412976 was only assembled into a partial sequence in a previous study [6]. However, the exact identification of viruses requires the complete genomes or even the full-length genomes. Using paired-end sequencing data, we reassembled these two virus genomes and obtained two full-length sequences to update EPI_ISL_412976 and EPI_ISL_412977 (**Supplementary 1**). Using 5’ UTR barcodes (**Introduction**), the betacoronaviruses RmYN01 and RmYN02 were identified as belonging to subgroup B. Using the InDels at M1 to M6, RmYN01 was further identified as belonging to the second class, respectively. Using the InDels at M1 to M4 and M6, RmYN02 was further identified as belonging to the first class but a recombinant SARS2-like CoV strain. RmYN02 was supposed to have a 3-nt deletion at the M5 site; however, it did not (**Table 1**). This led us to report—for the first time—a recombination event in *ORF8* at the whole-gene level in a bat, which had been co-infected by two betacoronavirus strains.

*ORF8* (**Figure 2**), existing only in betacoronavirus subgroup B, was considered to have played a significant role in adaptation to human hosts following interspecies transmission [7] via the modification of viral replication [8]. A 29-nt deletion in SARS-CoV (GenBank: AY274119) was reported and considered to be associated with attenuation during the early stage of human-to-human transmission [8].

A 382-nt deletion during the early evolution of SARS-CoV-2 (GISAID: EPI_ISL_414378) was also reported [12], but associated with attenuation without changes in its replication [13]. Although many recombination events in *ORF8* of betacoronaviruses have been reported in sequence analysis results, it is difficult to determine whether they were recombination events or small-size mutation (InDel & SNP) accumulation as most of them only occurred over very small genomic regions, excepting a few events (e.g., the 382-nt deletion in *ORF8* [12]). In the present study, the discovery of a recombination event in *ORF8* at the whole-gene level led to the determination of three types (**see below**) of *ORF8* genes in the betacoronavirus subgroup B, providing clues to understand the functions of *ORF8*.

### Conjoint analysis of phylogeny and molecular function

Based on conjoint analysis of phylogeny and molecular function that was proposed in our previous study [9], genes (i.e. *ORF1a, S1* and *ORF8*) containing the recombination regions under high selection pressure must be removed in phylogenetic analysis. Using large segments (**Supplementary 1**) spanning *S2*, *ORF3a, 3b*, envelope (*E*), *M, ORF6, 7a, 7b, N*(*9a*) and *ORF10* (**Table 1**), phylogenetic tree 1 (**Figure 3A**) showed that 19 of 21 betacoronaviruses from subgroup B (**Materials and Methods**) were classified into two major clades, corresponding to the first and second class classified using the InDels at six sites, respectively: (1) the first major clade, named the SARS-CoV-2 cluster, includes SARS-CoV-2 and all SARS2-like CoVs (from bats and pangolins); and (2) the second major clade includes two clusters—the SARS-CoV cluster including SARS-CoV and a few the most closely related SARS-like CoVs (from bats or civets) and the SARS-like CoV cluster including all other SARS-like CoVs. Therefore, these 19 betacoronaviruses were classified into the SARS-CoV-2, SARS-CoV and SARS-like CoV clusters named clusters 1, 2 and 3 (**Figure 3A**), respectively, while the other two betacoronaviruses (i.e. WIV1 and RmYN02) are recombinant strains. With more data from SARS2-like CoVs available, the SARS-CoV-2 cluster (a temporary result) need be further divided into the SARS-CoV-2 and SARS2-like CoV clusters by the presence of junction FCS “RRA**<u>R</u>**” (**see below**), while the SARS-CoV class was divided into the SARS-CoV and SARS-like CoV clusters by the types of *ORF8*.

**Figure 3.**
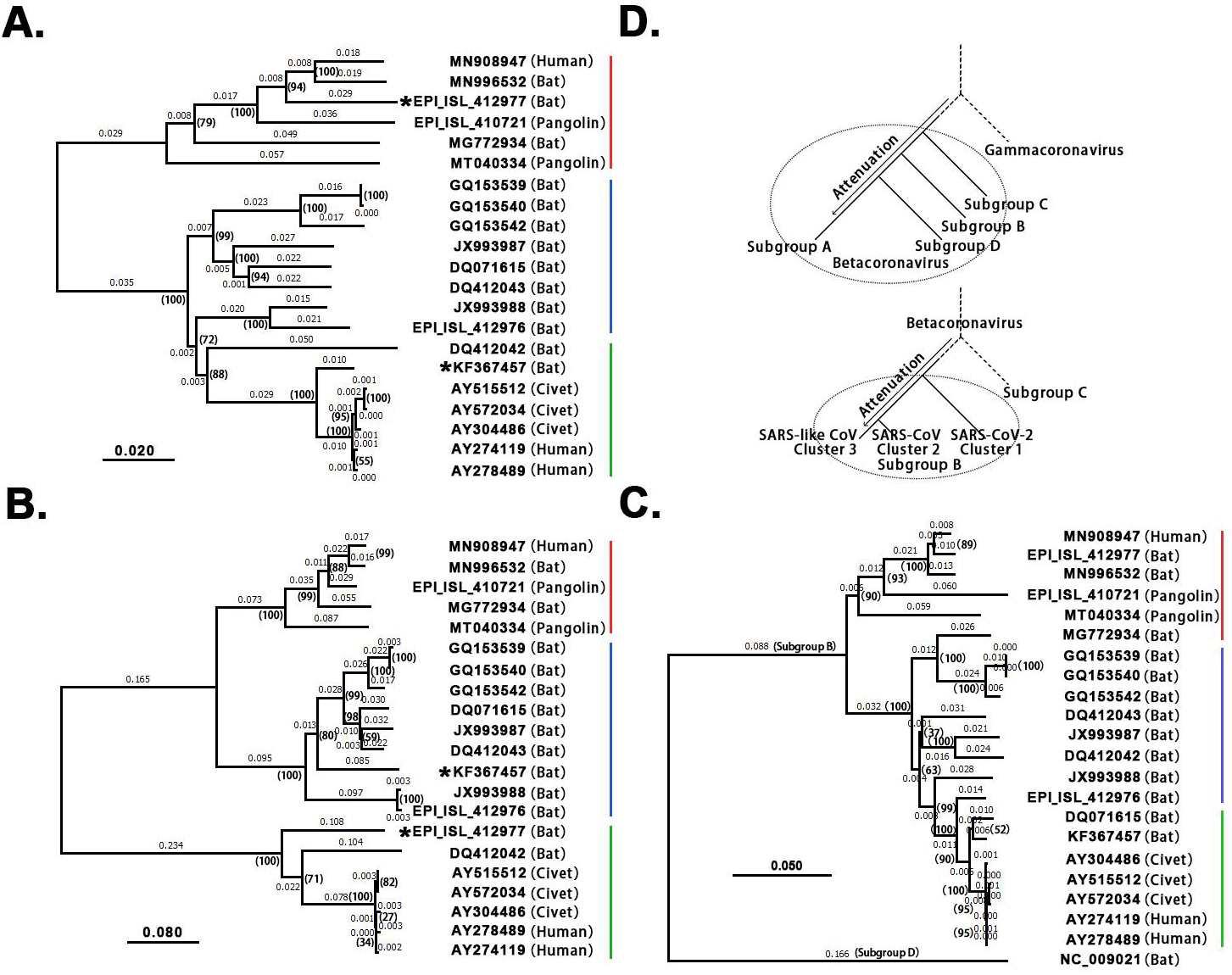
Phylogenetic analysis and evolution of betacoronavirus.

Using only *ORF8* (**Supplementary 1**), phylogenetic tree 2 (**Figure 3B**) also showed that the 19 betacoronaviruses were classified into the SARS-CoV-2, SARS-CoV and SARS-like CoV clusters. However, this tree did not reflect the evolutionary relationship of the three clusters due to the recombination events of *ORF8*. Using 21 CDSs of *nsp12* (RNA-dependent RNA polymerase, RdRP), the rooted phylogenetic tree 3 (**Figure 3C**) was construct to confirm the evolutionary relationship of the three clusters in tree 1 (**Figure 3A**).In phylogenetic tree 2, the SARS-CoV-2, SARS-CoV and SARS-like CoV clusters have types 1, 2 and 3 ORF8 genes, respectively. Type 1 *ORF8* genes possess low nucleotide identities (below 70%) to type 3 *ORF8* genes, while type 2 *ORF8* genes are so highly divergent from types 1 and 3 *ORF8* genes, they cannot be well aligned to calculate nucleotide identities between type 2 *ORF8* genes and types 1 or 3 *ORF8* genes. As RmYN02 belongs to the cluster 1 (**Figure 3A**) but has a type 2 rather than type 1 *ORF8* (**Figure 3B**), RmYN02 is a recombinant SARS2-like CoV strain. This discovery indicated that recombination occurred across the SARS-CoV-2 and SARS-CoV clusters, which has potential to generate a new strain more dangerous than SARS-CoV-2 and SARS-CoV.

Comparing phylogenetic tree 1 (**Figure 3A**) using large segments with 2 (**Figure 3B**) using only *ORF8* genes, all betacoronaviruses were consistently classified into the same clusters in both trees, except RmYN02 and the SARS-like CoV strain WIV1 (GenBank: KF367457).

The accession numbers of the GenBank or GISAID databases were used to represent the viral genomes: MN908947: SARS-CoV-2; MN996532: the SARS2-like CoV strain RaTG13; EPI_ISL_412977: the SARS2-like CoV strain RmYN02; EPI_ISL_412976: the SARS-like CoV strain RmYN01; KF367457: the SARS-like CoV strain WIV1; AY274119: the SARS-CoV strain Tor2; AY278489: the SARS-CoV strain GD01. Decimal above the branches are phylogenetic distances calculated using the NJ method with a bootstrap test (1000 replicates). The bootstrap values (marked by parentheses) were in the format for displaying percentages with “%” omitted. 19 of 21 betacoronaviruses from subgroup B were classified into the SARS-CoV-2 (red), SARS-like CoV (blue) and SARS-CoV (green) clusters, while the other two betacoronaviruses (i.e. WIV1 and RmYN02) are recombinant strains. **A**. Phylogenetic tree 1 was built using large segments spanning *S2, ORF3a*, 3b, *E*, *M, ORF6, 7a, 7b, N(9b)* and *ORF10* (**Table 1**). **B**. Phylogenetic tree 2 was built using *ORF8*. As type 2 *ORF8* genes cannot be well aligned to types 1 or 3 *ORF8* genes to calculate nucleotide identities, the distances between the SARS-CoV cluster and the SARS-CoV-2 or SARS-like CoV clusters are not accurate. **C**. Phylogenetic tree 3 was built using CDSs of *nsp12* (RNA-dependent RNA polymerase, RdRP). HKU9-CoV (RefSeq: NC_009021) from subgroup D was used as an outgroup species. **D**. MERS-CoV (GenBank: JX869059), SARS-CoV-2 (GenBank: MN908947), HKU9-CoV (RefSeq: NC_009021), MHV (RefSeq: NC_001846) and IBV (RefSeq: NC_001451) were used to represent betacoronavirus subgroup C, B, D, A and gammacoronavirus, respectively in the upper phylogenetic tree; SARS-CoV-2 (GenBank: MN908947), SARS-CoV (GenBank: AY278489), RmYN01 (GISAID: EPI_ISL_412976) and MERS-CoV (GenBank: JX869059) were used to represent the SARS-CoV-2, SARS-CoV and SARS-like CoV clusters from betacoronavirus subgroup B and betacoronavirus subgroup C, respectively in the lower phylogenetic tree.

WIV1 was classified into cluster 2, but has a type 3 rather than type 2 *ORF8*, WIV1 is a recombinant SARS-like CoV strain. WIV1, isolated from Chinese horseshoe bats (*Rhinolophus sinicus*), was considered the most closely related to SARS-CoV but not its immediate ancestor [14]. A previous study predicted the immediate ancestor of SARS-CoV based on the following hypothesis: the ancestor of SARS-like CoVs from civets was a recombinant virus with *ORF8* originating from greater horseshoe bats (*Rhinolophus ferrumequinum*) and other genomic regions originating from different horseshoe bats [7]. However, whether these recombination events occurred in bats or civets remains unclear [7]. Both phylogenetic tree 1 (**Figure 3A**) and 2 (**Figure 3B**) consistently revealed that SARS-CoV-2 is most closely related to the well-known strain RaTG13 (GenBank: MN996532) isolated from intermediate horseshoe bats (*Rhinolophus affinis*). However, RaTG13 is unlikely to be the immediate ancestor of SARS-CoV-2 due to lack of the junction FCS “RRA**<u>R</u>**”. In addition, all pangolin (*Manis javanica*) betacoronaviruses investigated using their public genomes (**Materials and Methods**) were identified as belonging to the SARS-CoV-2 cluster. However, further analysis of these genomes does not support that pangolins are the intermediate host(s) of SARS-CoV-2 [15] for three main reasons: (1) pangolin betacoronaviruses do not have the junction FCS “RRA**<u>R</u>**” [3]; (2) the strains (e.g. GISAID: EPI_ISL_410721) are farther from SARS-CoV-2 than RaTG13 in the phylogenetic tree 1 (**Figure 3A**); and (3) pangolin betacoronaviruses are unlikely to lose “RRA**<u>R</u>**” so soon, as betacoronavirus subgroup A lost a junction FCS after a long-term evolutionary change. This suggested that the intermediate host(s) of SARS-CoV-2 carry at least one strain of CoV with junction FCS “RRA**<u>R</u>**” in the S protein. WIV1, RaTG13, RmYN01, RmYN02 and pangolin betacoronaviruses are attenuated variants of SARS-CoV-2 and SARS-CoV (**see below**).

### Determination of *ORF8*’ functions

Next, we conducted further research on the biological functions of *ORF8*. RmYN01 and RmYN02 were simultaneously detected in a bat, providing a special opportunity to compare their copy numbers. As RmYN01 and RmYN02 have type 3 and type 2 *ORF8* genes, respectively, the difference between the copy numbers of RmYN01 and that of RmYN02 can be estimated by their relative RNA abundances to test a previous hypothesis that type 2 *ORF8* genes increase replication efficiency of viruses. Aligning RNA-seq data to the genomes of RmYN01 and RmYN02, our calculation showed that the RmYN01 genome was covered 99.85% of its length with an average depth of 32.89 (**Figure 2A**), while the RmYN02 genome was covered 99.89% with an average depth of 298.99 (**Figure 2B**). The RNA abundance of RmYN02 is 9 times that of RmYN01. Based on the “leader-to-body fusion” model explaining the replication and transcription of CoVs [5], the difference in RNA abundance of the *ORF1a* and *ORF1b* genes (**Figure 2AB**) resulted from CoV replication, rather than transcription. Therefore, this result suggests that type 2 *ORF8* (named enhanced *ORF8*) genes increase replication efficiency of RmYN02, ruling out the possibility that transcription contributes to the difference in RNA abundance of the two virus strains. Our study ultimately determined that *ORF8* enhances the viral replication.

### Evolution and outbreak of betacoronavirus

Receptor binding abilities, junction FCSs, strong first RBSs and enhanced *ORF8*s (**see above**) are main factors contributing to transmission, virulence and host adaptability of CoVs. By analysis of these main factors in betacoronavirus genomes (**Materials and Methods**), we concluded: (1) rapid recombination of viral genomes provides CoV the strong ability of cross-species transmission and outbreak; (2) the immediate ancestor of betacoronavirus was most likely to have two junction FCS and a strong first RBS, and it transmitted across species during its outbreak; (3) after a period of adaption in new hosts, betacoronavirus was attenuated to spread widely and persist in the host population by loss of abilities attributed to one or more factors (e.g. junction FCSs); and (4) the strong recombination ability of CoVs integrated these factors to generate multiple recombinant strains, very a few of which evolved into super virus strains (e.g. SARS-CoV and SARS-CoV-2) causing pandemics by nature selection.

In the betacoronavirus subgroup C (**Figure 4A**), middle east respiratory syndrome coronavirus (MERS-CoV) has two junction FCSs. The first one “RST**<u>R</u>**”, located at position 694 in the S protein (noted as R694), is nonfunctioning, as a result of attenuation, because there is a disulfide bond cross the junction FCS R694. However, the second junction FCS “RSV**<u>R</u>**” (R751) is still functional. Originated from the same ancestor of MERS-CoV, MERS-like CoVs (e.g. hedgehog CoV) were further attenuated by loss of two junction FCSs. In the betacoronavirus subgroup B (**Figure 4AB**), SARS-CoV-2 (GenBank: MN908947) has the junction FCS “RRA**<u>R</u>**” (R685) and lost a junction FCS by substituting “KNTQ” (R779) for “RNT**<u>R</u>**”, as a result of attenuation. All SARS-2 like CoVs (from bats or pangolins [3]) in the SARS-CoV-2 cluster were further attenuated by loss of “RRA**<u>R</u>**” and substituting “KNTQ” for “RNT**<u>R</u>**”. The immediate ancestor of SARS-CoV was an attenuated variant of SARS-CoV-2 by loss of “RRA**<u>R</u>**” (e.g R667) and with inaccessible “RNT**<u>R</u>**” (R761) which has secondary structures in helix rather than coil. All SARS-like CoVs in the SARS-like CoV cluster were further attenuated by loss of “RRA**<u>R</u>**” and substituting “KNTQ” for “RNT**<u>R</u>**”. In the betacoronavirus subgroup D, HKU9-CoV was attenuated by loss of two junction FCSs but still have a strong first RBS. In the betacoronavirus subgroup A (**Figure 4A**), although almost all strains (e.g. HCoV-OC43 and HCoV-HKU1) still have one junction FCS, they do not have strong first RBSs and enhanced *ORF8*s. These strains were heavily attenuated from the immediate ancestor of betacoronavirus due to complex reasons. Firstly, the average arginine (R) pencentage of S proteins from betacoronaviruses of the subgroup A except mouse hepatitis virus (MHV) 2.63% is significantly lower than those from MHV and betacoronaviruses of the subgroup B, C and D (3.34%, 3.33%, 3.32% and 3.33%). This indicated that accumulated mutations caused attenuation by loss of arginine residues, since arginine residues are indispensable for the protease cleavage sites. Other reasons may include the loss of strong first RBSs and genetic events in the transcription regulatory sequences [5], an important factor that is not further investigated in the present study, but merit further investigation in the future.

**Figure 4.**
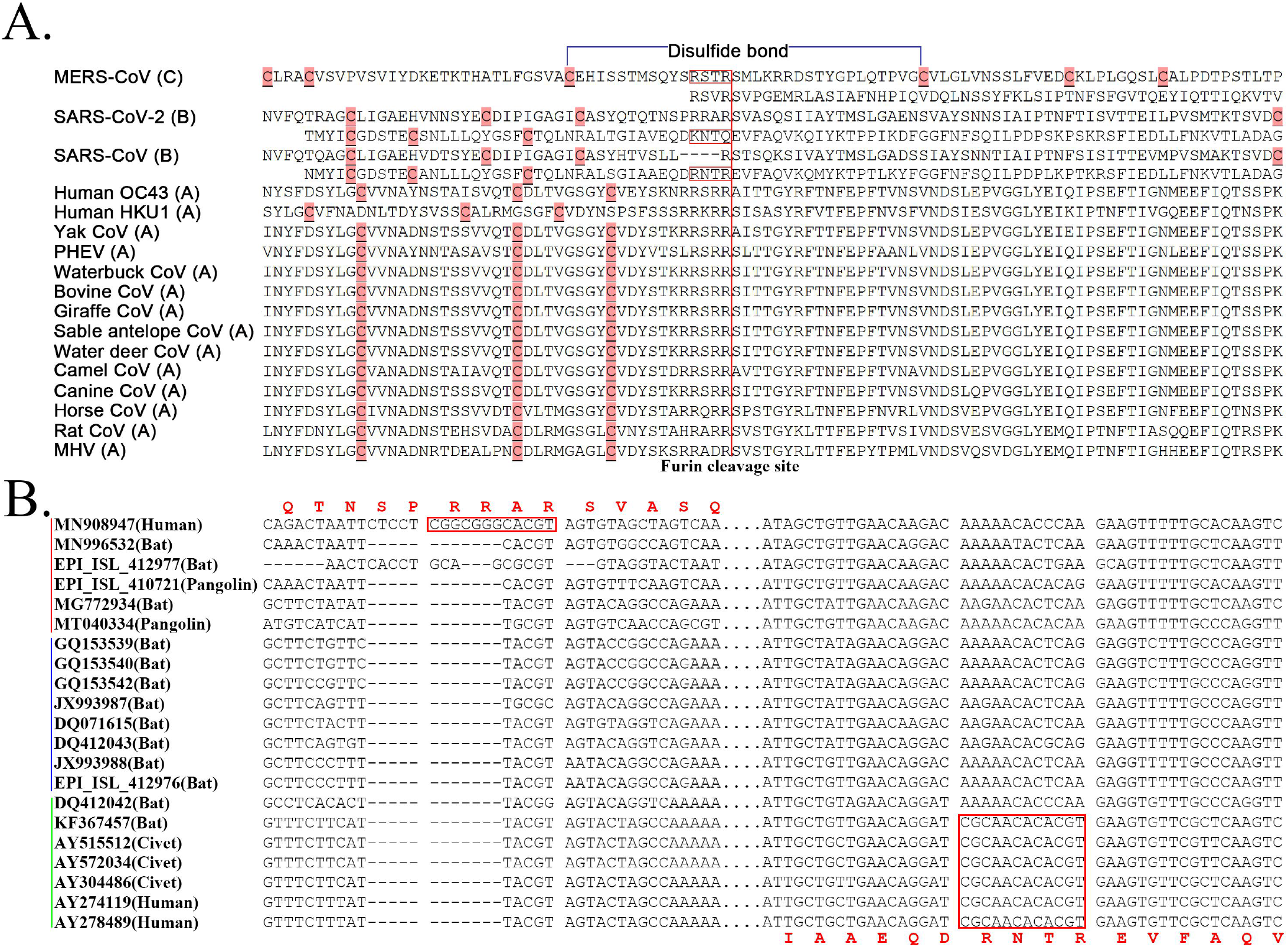
Junction furin cleavage sites of betacoronaviruses. **A**. Two regions having potential to contain junction furin cleavage sites (FCSs) are showed for MERS-CoV, SARS-CoV-2 and SARS-CoV, while only one region is showed for other CoVs. Junction FCSs (in red box) are non-functioning, lost or inaccessible due to different reasons. The disulfide bond (in blue color) is only cross “RSTR” of MERS-CoV. MERS-CoV (GenBank: JX869059) belongs to subgroup C; SARS-CoV (GenBank: AY278489) and SARS-CoV-2 (GenBank: MN908947) belong to subgroup B; Human OC43 (GenBank: KF530084), Human HKU1 (GenBank: KF686346), Yak CoV (GenBank: MH810163), PHEV (GenBank: KY419107), Waterbuck CoV (GenBank: FJ425186), Bovine CoV (GenBank: MH043954), Giraffe CoV (GenBank: EF424622), Sable antelope CoV (GenBank: EF424621), Water deer CoV (GenBank: MG518518), Camel CoV (GenBank: MN514963), Canine CoV (GenBank: JX860640), Horse CoV (GenBank: LC061274), Rat CoV (GenBank: JF792617) and MHV (GenBank: AF029248) belong to subgroup A. PHEV: porcine hemagglutinating encephalomyelitis virus; MHV: mouse hepatitis virus. **B**. Two regions having potential to contain junction FCSs (in red box) are showed at the nucleotide level. The SARS-CoV-2, SARS-like CoV and SARS-CoV clusters were marked by red, blue and green lines.

Guided by conjoint analysis of molecular function and phylogeny, we concluded the following (**Figure 3D**): (1) in general, betacoronaviruses (and even CoVs) were and are undergoing attenuation to spread widely and persist in host population after every outbreak; (2) the immediate ancestor of subgroup C (e.g. MER-CoV) was the most closely related to the immediate ancestor of betacoronavirus with slight attenuation; (3) the immediate ancestors of subgroup B and D diverged subsequently and were further attenuated; and (4) betacoronaviruses from the subgroup A were the most heavily attenuated and have the highest diversity in their genomes and hosts. In the same principle (**Figure 3D**), (1) the immediate ancestor of the SARS-CoV-2 cluster was the most closely related to the immediate ancestor of the subgroup B with slight attenuation; (2) the immediate ancestor of the SARS-CoV cluster diverged subsequently and was further attenuated; and (3) the SARS-like CoV cluster was the most heavily attenuated and has the highest diversity in the genomes and hosts. All the SARS-like CoVs (e.g. WIV1 and RmYN01) in the SARS-like CoV cluster are attenuated variants of SARS-CoV, while the SARS2-like CoVs (e.g. RaTG13, RmYN02 and betacoronaviruses from pangolins) in the SARS-CoV-2 cluster are attenuated variants of SARS-CoV-2. As recombinant betacoronavirus, the immediate ancestor of SARS-CoV-2 is characterized by the junction FCS “RRA**<u>R</u>**”, while the immediate ancestor of SARS-CoV is characterized by the enhanced *ORF8*. Therefore, WIV1 without the enhanced *ORF8* and RaTG13 without the junction FCS “RRA**<u>R</u>**” may contribute to, but are not the immediate ancestors of SARS-CoV and SARS-CoV-2, respectively.

## Conclusions

The outbreaks of MER-CoV, SARS-CoV and SARS-CoV-2 were triggered by recombination events, not accumulated mutations. So it is not suitable to estimate their divergence time using current theories in evolutionary biology. The origins of the junction FCS “RRA**<u>R</u>**” and the enhanced *ORF8* are still unknown. Therefore, the recombinant strains (e.g. WIV1 and RmYN02) are identified based on the reference strains that were usually reported before the recombinant strains. This reflects the phylogenetic relationship between them, not the actual recombinant events which occurred to generate the recombinant strains. Future investigation need be conducted to search for the betacoronavirus strains that provided the junction FCS “RRA**<u>R</u>**” and the enhanced *ORF8* to SARS-CoV-2 and SARS-CoV, respectively. our studies also suggest rthat the nucleotide sequences of the junction FCS “RRA**<u>R</u>**” and *ORF8* may originate from non-viral species.

Receptor binding abilities, junction FCSs, strong first RBSs and enhanced *ORF8*s are main factors contributing to transmission, virulence and host adaptability of CoVs. Junction FCSs and enhanced *ORF8*s increase the efficiencies in viral entry into cells and replication, respectively while strong first RBSs enhance the translational initiation. The strong recombination ability of CoVs integrated these factors to generate multiple recombinant strains, two of which evolved into SARS-CoV and SARS-CoV-2 by nature selection, resulting in the SARS and COVID-19 pandemics. Based on our theories, two predictions can be made: (1) more attenuated (by loss of Junction FCSs or enhanced *ORF8*s) variants of SARS-CoV-2 will be reported; and (2) SARS2-like CoV with at least one junction FCS will be eventually detected in bats.

## Materials and Methods

The software VirusDetect [16] was used to detect viruses in RNA-seq data from [6]. The software Fastq_clean [17] was used for RNA-seq data cleaning and quality control. The genomes of RmYN01 and RmYN02 (GISAID: EPI_ISL_412976 and EPI_ISL_412977) were reassembled by aligning RNA-seq data on two closest reference genomes JX993988 and MN908947. SVDetect v0.8b and SVFilter [18] were used to removed abnormal aligned reads. Several haploid contigs (**Supplementary 1**) highly similar to the complete RmYN01 genome were also assembled. This suggested that there exists more than one betacoronavirus strain belonging to the SARS-like CoV cluster in the same sample, from which RmYN01 and RmYN02 were detected.

1,265 genome sequences of betacoronaviruses (in group A, B, C and D) were downloaded from the NCBI Virus database (https://www.ncbi.nlm.nih.gov/labs/virus) in our previous study [3]. Among these genomes, 292 belongs to betacoronavirus subgroup B. 35 genomes of betacoronavirus subgroup B were also downloaded from the GISAID database. In our previous study, 10 complete genomes of betacoronavirus subgroup B (GenBank: JX993987, JX993988, GQ153539, GQ153540, GQ153542, DQ071615, DQ412043, AY515512, AY572034 and DQ497008) were selected and used for the analysis [9]. To trace the origin of SARS-CoV, five complete genomes were added in the present study. They are DQ412042 (SARS-like CoV from *Rhinolophus ferrumequinum*), AY274119 (SARS-like CoV from human in Toronto, Tor2 [19]), AY278489 (SARS-like CoV from human in Guangdong, GD01 [20]), AY304486 (SARS-like CoV from civet [21]) and KF367457 (SARS-like CoV from bat [22]). DQ497008 was removed as a redundant sequence of AY274119 and AY278489. To trace the origin of SARS-CoV-2, three complete genomes were added. They are MN908947 (SARS-CoV-2), MN996532 (SARS2-like CoV hosted in Intermediate Horseshoe bats (*Rhinolophus affinis*) from Yunnan) and MG772934 (SARS2-like CoV hosted in Chinese horseshoe bats (*Rhinolophus sinicus*) from Zhejiang). A SARS2-like CoV (GISAID: EPI_ISL_410721) from pangolins (Collected in Guangdong, China) and a SARS2-like CoV (GenBank: MT040334) from pangolins (Collected in Guangxi, China) were also used. Plus RmYN01 and RmYN02 (GISAID: EPI_ISL_412976 and EPI_ISL_412977), totally 21 complete genomes were used for the phylogenetic analysis applying the neighbour joining (NJ) method. Sequence alignment was performed using the Bowtie v0.12.7 software with paired-end alignment allowing 3 mismatches; mutation detection and other data processing were carried out using Perl scripts; the phylogenetic analysis was performed using MEGA v7.0.26; Statistics and plotting were conducted using the software R v2.15.3 the Bioconductor packages [23]. The structure of S (**Supplementary 2**) was predicted using trRosetta [24].

## Acknowledgments

First, we thank Professor Weifeng Shi from Shandong First Medical University for his RNA-seq data sharing. We are grateful for the help from the following faculty members of College of Life Sciences at Nankai University: Xuetao Cao, Deling Kong, Quan Chen, Wenjun Bu, Ting Ma, Tao Zhang, Dawei Huang, Mingqiang Qiao, Yanqiang Liu, Bingjun He and Zhen Ye. We also appreciate the cooperation and support from Professor Ze Chen from Hebei Normal University. This project was supported by Yunnan Provincial Department of Education Science Research Fund Project Funding (No. 2018JS188) to Shunmei Chen and Tianjin Key Research and Development Program of China (19YFZCSY00500) to Shan Gao. We would like to thank Editage (www.editage.cn) for English language translation. This manuscript was online as a preprint on July 22^nd^, 2020 at https://biorxiv.org/cgi/content/short/2020.07.22.213926v1.

## Author contributions statements

SG conceived the project. SG and GD supervised this study. XJ and SC conducted programming. XL, LW, and TY downloaded, managed and processed the data. JY predicted the structure of the S protein. JR analyzed the structure of S1. SG drafted the main manuscript text. SG and ZH revised the manuscript.

## REFERENCES

1. R.L. Graham and R.S. Baric, Recombination, Reservoirs, and the Modular Spike: Mechanisms of Coronavirus Cross-Species Transmission. Journal of Virology, 2010. 84(7): p. 3134–3146.

2. C. Jiayuan, J. Shi, O. Yau Tung, C. Liu, X. Li, Q. Zhao, R. Jishou and G. Shan, Bioinformatics Analysis of the 2019 Novel Coronavirus Genome. Chinese Journal of Bioinformatics (In Chinese), 2020. 18(2): p. 96–102.

3. X. Li, G. Duan, W. Zhang, J. Shi, J. Chen, S. Chen, S. Gao and J. Ruan, A Furin Cleavage Site Was Discovered in the S Protein of the 2019 Novel Coronavirus. Chinese Journal of Bioinformatics (In Chinese), 2020. 18(2): p. 103–108.

4. G. Duan, J. Shi, Y. Xuan, J. Chen, C. Liu, J. Ruan, S. Gao and X. Li, 5’ UTR Barcode of the 2019 Novel Coronavirus Leads to Insights into Its Virulence. Chinese Journal of Virology (In Chinese), 2020. 36(3): p. 365–369.

5. D. Kim, J.-Y. Lee, J.-S. Yang, J.W. Kim, V.N. Kim and H. Chang, The Architecture of SARS-CoV-2 Transcriptome. Cell, 2020. 181(4): p. 914–921.

6. H. Zhou, X. Chen, T. Hu, J. Li, H. Song, Y. Liu, P. Wang, D. Liu, J. Yang and E.C. Holmes, A novel bat coronavirus closely related to SARS-CoV-2 contains natural insertions at the S1/S2 cleavage site of the spike protein. Current Biology, 2020.

7. S.K.P. Lau, Y. Feng, H. Chen, H.K.H. Luk, W.H. Yang, K.S.M. Li, Y.Z. Zhang, Y. Huang, Z.Z. Song and W.N. Chow, SARS coronavirus ORF8 protein is acquired from SARS-related coronavirus from greater horseshoe bats through recombination. Journal of Virology, 2015: p. JVI.01048-15.

8. D. Muth, V.M. Corman, H. Roth, T. Binger, R. Dijkman, L.T. Gottula, F. Gloza-Rausch, A. Balboni, M. Battilani and D. Rihtarič, Attenuation of replication by a 29 nucleotide deletion in SARS-coronavirus acquired during the early stages of human-to-human transmission. Scientific Reports, 2018. 8(1).

9. C. Liu, Z. Chen, Y. Hu, H. Ji, D. Yu, W. Shen, S. Li, J. Ruan, W. Bu and S. Gao, Complemented Palindromic Small RNAs First Discovered from SARS Coronavirus. Genes, 2018. 9(9): p. 1–11.

10. D. Ellinghaus, F. Degenhardt, L. Bujanda, M. Buti, A. Albillos, P. Invernizzi, J. Fernández, D. Prati, G. Baselli, R. Asselta, M.M. Grimsrud, C. Milani, F. Aziz, J. Kässens, S. May, M. Wendorff, L. Wienbrandt, F. Uellendahl-Werth, T. Zheng, X. Yi, R. de Pablo, A.G. Chercoles, A. Palom, A.-E. Garcia-Fernandez, F. Rodriguez-Frias, A. Zanella, A. Bandera, A. Protti, A. Aghemo, A. Lleo, A. Biondi, A. Caballero-Garralda, A. Gori, A. Tanck, A. Carreras Nolla, A. Latiano, A.L. Fracanzani, A. Peschuck, A. Julià, A. Pesenti, A. Voza, D. Jiménez, B. Mateos, B. Nafria Jimenez, C. Quereda, C. Paccapelo, C. Gassner, C. Angelini, C. Cea, A. Solier, D. Pestaña, E. Muñiz-Diaz, E. Sandoval, E.M. Paraboschi, E. Navas, F. García Sánchez, F. Ceriotti, F. Martinelli-Boneschi, F. Peyvandi, F. Blasi, L. Téllez, A. Blanco-Grau, G. Hemmrich-Stanisak, G. Grasselli, G. Costantino, G. Cardamone, G. Foti, S. Aneli, H. Kurihara, H. ElAbd, I. My, I. Galván-Femenia, J. Martín, J. Erdmann, J. Ferrusquía-Acosta, K. Garcia-Etxebarria, L. Izquierdo-Sanchez, L.R. Bettini, L. Sumoy, L. Terranova, L. Moreira, L. Santoro, L. Scudeller, F. Mesonero, L. Roade, M.C. Rühlemann, M. Schaefer, M. Carrabba, M. Riveiro-Barciela, M.E. Figuera Basso, M.G. Valsecchi, M. Hernandez-Tejero, M. Acosta-Herrera, M. D’Angiò, M. Baldini, M. Cazzaniga, M. Schulzky, M. Cecconi, M. Wittig, M. Ciccarelli, M. Rodríguez-Gandía, M. Bocciolone, M. Miozzo, N. Montano, N. Braun, N. Sacchi, N. Martínez, O. Özer, O. Palmieri, P. Faverio, P. Preatoni, P. Bonfanti, P. Omodei, P. Tentorio, P. Castro, P.M. Rodrigues, A. Blandino Ortiz, R. de Cid, R. Ferrer, R. Gualtierotti, R. Nieto, S. Goerg, S. Badalamenti, S. Marsal, G. Matullo, S. Pelusi, S. Juzenas, S. Aliberti, V. Monzani, V. Moreno, T. Wesse, T.L. Lenz, T. Pumarola, V. Rimoldi, S. Bosari, W. Albrecht, W. Peter, M. Romero-Gómez, M. D’ Amato, S. Duga, J.M. Banales, J.R. Hov, T. Folseraas, L. Valenti, A. Franke and T.H. Karlsen, Genomewide Association Study of Severe Covid-19 with Respiratory Failure. New England Journal of Medicine, 2020.

11. M.P. Lins and S. Smaniotto, Potential impact of SARS-CoV-2 infection on the thymus. Canadian Journal of Microbiology, 2020. 66(8): p. 1–20.

12. Y.C. Su, D.E. Anderson, B.E. Young, F. Zhu, M. Linster, S. Kalimuddin, J.G. Low, Z. Yan, J. Jayakumar, L. Sun, G.Z. Yan, I.H. Mendenhall, Y.-S. Leo, D.C. Lye, L.-F. Wang and G.J. Smith, Discovery of a 382-nt deletion during the early evolution of SARS-CoV-2. bioRxiv, 2020. 1(1): p. 1–23.

13. B.E. Young, S.-W. Fong, Y.-H. Chan, T.-M. Mak, L.W. Ang, D.E. Anderson, C.Y.-P. Lee, S.N. Amrun, B. Lee, Y.S. Goh, Y.C.F. Su, W.E. Wei, S. Kalimuddin, L.Y.A. Chai, S. Pada, S.Y. Tan, L. Sun, P. Parthasarathy, Y.Y.C. Chen, T. Barkham, R.T.P. Lin, S. Maurer-Stroh, Y.-S. Leo, L.-F. Wang, L. Renia, V.J. Lee, G.J.D. Smith, D.C. Lye and L.F.P. Ng, Effects of a major deletion in the SARS-CoV-2 genome on the severity of infection and the inflammatory response: an observational cohort study. The Lancet, 2020. 396(10248): p. 1–9.

14. G. Xing-Yi, L. Jia-Lu, Y. Xing-Lou, C. Aleksei A, Z. Guangjian, E. Jonathan H, M. Jonna K, H. Ben, Z. Wei, P. Cheng, Z. Yu-Ji, L. Chu-Ming, T. Bing, W. Ning, Z. Yan, C. Gary, Z. Shu-Yi, W. Lin-Fa, D. Peter and S. Zheng-Li, Isolation and characterization of a bat SARS-like coronavirus that uses the ACE2 receptor. Nature, 2013. 503(7477): p. 535–538.

15. T.T.-Y. Lam, N. Jia, Y.-W. Zhang, M.H.-H. Shum, J.-F. Jiang, H.-C. Zhu, Y.-G. Tong, Y.-X. Shi, X.-B. Ni, Y.-S. Liao, W.-J. Li, B.-G. Jiang, W. Wei, T.-T. Yuan, K. Zheng, X.-M. Cui, J. Li, G.-Q. Pei, X. Qiang, W.Y.-M. Cheung, L.-F. Li, F.-F. Sun, S. Qin, J.-C. Huang, G.M. Leung, E.C. Holmes, Y.-L. Hu, Y. Guan and W.-C. Cao, Identifying SARS-CoV-2-related coronaviruses in Malayan pangolins. Nature, 2020. 583(7815): p. 282–285.

16. Y. Zheng, S. Gao, C. Padmanabhan, R. Li, M. Galvez, D. Gutierrez, S. Fuentes, K. Ling, J.F. Kreuze and Z. Fei, VirusDetect: An automated pipeline for efficient virus discovery using deep sequencing of small RNAs. Virology, 2016. 500(2017): p. 130–138.

17. M. Zhang, F. Zhan, H. Sun, X. Gong, Z. Fei and S. Gao. Fastq_clean: An optimized pipeline to clean the Illumina sequencing data with quality control. in Bioinformatics and Biomedicine (BIBM), 2014 IEEE International Conference on. 2014. IEEE.

18. F. Zhang, T. Xu, L. Mao, S. Yan, X. Chen, Z. Wu, R. Chen, X. Luo, J. Xie and S. Gao, Genomewide analysis of Dongxiang wild rice (Oryza rufipogon Griff.) to investigate lost/acquired genes during rice domestication. BMC plant biology, 2016. 16(1): p. 1–11.

19. R. He, F. Dobie, M. Ballantine, A. Leeson, Y. Li, N. Bastien, T. Cutts, A. Andonov, J. Cao and T.F. Booth, Analysis of multimerization of the SARS coronavirus nucleocapsid protein. Biochemical and biophysical research communications, 2004. 316(2): p. 476–483.

20. Q. Wu, Y. Zhang, H. Lü, J. Wang, X. He, Y. Liu, C. Ye, W. Lin, J. Hu and J. Ji, The E protein is a multifunctional membrane protein of SARS-CoV. Genomics, proteomics & bioinformatics, 2003. 1(2): p. 131–144.

21. Y. Guan, B. Zheng, Y. He, X. Liu, Z. Zhuang, C. Cheung, S. Luo, P. Li, L. Zhang and Y. Guan, Isolation and characterization of viruses related to the SARS coronavirus from animals in southern China. Science, 2003. 302(5643): p. 276–278.

22. X.-Y. Ge, J.-L. Li, X.-L. Yang, A.A. Chmura, G. Zhu, J.H. Epstein, J.K. Mazet, B. Hu, W. Zhang and C. Peng, Isolation and characterization of a bat SARS-like coronavirus that uses the ACE2 receptor. Nature, 2013. 503(7477): p. 535–538.

23. S. Gao, J. Ou and K. Xiao, R language and Bioconductor in bioinformatics applications(Chinese Edition). 2014, Tianjin: Tianjin Science and Technology Translation Publishing Ltd.

24. J. Yang, I. Anishchenko, H. Park, Z. Peng and D. Baker, Improved protein structure prediction using predicted interresidue orientations. Proceedings of the National Academy of ences, 2020. 117(3): p. 201914677.

